# Novel orthotopic model of neuroblastoma in RNU homozygous rats

**DOI:** 10.1101/2023.01.16.523967

**Authors:** ReidAnn Sever, Lauren Taylor Rosenblum, Lori Schmidt, Sean Hartwick, Kristina Schwab, Yijen Wu, Itay Raphael, Miguel Reyes, Wilson Barry Edwards, Marcus Malek, Gary Kohanbash

## Abstract

Neuroblastoma, the most common extracranial solid malignancy in children, accounts for 15% of pediatric cancer deaths despite multimodal therapy including surgical resection. Unfortunately, complete surgical resection remains challenging due to encasement of major neurovascular structures, unclear tumor margins, and remote nodal disease. While mouse models of neuroblastoma are extremely valuable for studying tumor biology and medical treatments, the small size renders the mouse model insufficient to evaluate novel surgical therapy.

Here, we have developed a novel rat model of neuroblastoma to facilitate further development of surgical treatment. Human neuroblastoma cells (SK-N-BE(2)) were injected into the adrenal gland of RNU nude rats. They developed 2 cm xenograft tumors at 5 weeks which were easily identifiable on MRI imaging and on visual inspection. The rats began losing weight and neared end stage at 7 weeks, at which point surgical resection was attempted. While surgical resection was technically feasible, the rats were too frail to survive surgery at the late stage. The pathology of the tumors was consistent with neuroblastoma: small round blue cells with strong PHOX2B staining. Thus, we present a novel rat neuroblastoma model that can be used for development of surgical techniques, such as the use of intraoperative contrast agents.

## Introduction

Neuroblastoma, a malignancy of neural crest cell origin, is the most common solid extracranial tumor in children. (1,2) Despite developments in chemotherapy, immunotherapy, and radiation treatments, patients with high-risk disease only have a 5-year survival rate of only 40-50%.(3) Multimodal treatment includes surgical resection, though obstacles such as encasement of neurovascular structures, unclear tumor margins, and remote disease challenge complete resection. Completeness of resection, however, affects both outcomes and the impact of maintenance therapy, while up to 50% of patients experience complications from surgery.(4) Our group is developing an intraoperative contrast agent for neuroblastoma to improve the identification of tumor and delineation of tumor margins. However, current models of neuroblastoma are inadequate to assess the operative impact of this agent, as well as other operative developments. The need was present to create a new model of neuroblastoma, particularly to study surgical innovations.

While there are well established models for neuroblastoma in mice, one benefit of a surgical tracer is the ability to find tumors that are deep within an organ or surrounded by connective tissue, which cannot be studied in a mouse model. Rat models offer multiple advantages over mouse models of human diseases. Most obviously, their larger size (RNU rats with average weight of 175 g are about 8 times larger than Nu/j mice with an average weight of 20 g), enables more delicate operative interventions. A larger animal model would enable growth of larger tumors and better study of subtle changes in tumor size and heterogeneity within a tumor. The longer lifespan also enables easier detection of significant differences in lifespan and time to tumor recurrence. Furthermore, toxicology experiments are more commonly performed in rats.

There are limited rat models of neuroblastoma. Nilsson *et al* described subcutaneous flank injections of SH-SY5Y human neuroblastoma cells in RNU rats. (5) Other subcutaneous xenograft models include a LAN-1 model and an SK-N-AS human neuroblastoma. (6,7) The location of the xenograft in the subcutaneous tissue rather than the adrenal gland does not sufficiently replicate the anatomy of the disease for surgical studies and the tumor growth and response might be different in a different environment. Adrenal-based xenografts are important not only to more closely replicate the disease, but as a model for surgical resection of the tumor in its most common location.

Engler *et al* created NB xenografts with LAN-1 human neuroblastoma cell injections into the tail vein, left adrenal gland, or aorta of 4-weei-old RNU rats. (8) Tumors grew rapidly after left adrenal gland injections (25,132 +/- 1,570 mm^3^ in 5 w), with smaller adrenal tumors in the aorta-injected group (5,300 +/0 1,300 mm^3^) and tiny tumors after tail vein injection (42 +/- 12 mm^3^). Furthermore, in rats with intra- aortal injections, 50% had liver micrometastases and 60% had femur micrometastases (as seen with anti- GD2 staining). To our knowledge, however, that model was last described over 20 years ago and no intra-adrenal xenograft model has been created or used since. Furthermore, SK-N-BE(2) cells have become an established cell line for xenograft tumors in mice, and they were similarly derived from the bone marrow of a 2-year-old with disseminated disease who had undergone multiple courses of chemotherapy and radiation. (9,10)

Here we aimed to create a neuroblastoma model for the development of surgical techniques. We report a novel adrenal xenograft neuroblastoma model in rats that closely resembles human disease.

## Materials & Methods

### Cell lines

SK-N-BE (2) (ATCC: no. CRL-2271) cells were cultured in Dulbecco’s Modified Eagle Medium (Lonza, Cat# BW12614F), supplemented with heat-inactivated FBS (Hyclone, Cat# SH3091003), sodium pyruvate (Lonza, Cat# BW13-115E), Antibiotic-Antimycotic (Gibco, Cat#15-240-062), beta-mercaptoethanol (Gibco, Cat#21-985-023), MEM NEAA (Gibco, Cat#11140-050), and 50 mg/mL Normicin (Invivogen, Cat# ant-nr-2) and maintained in a 37°C humidified incubator with 5% CO_2_.

### Animal studies

Animal studies were performed in accordance with the protocols approved by the Institutional Animal Care and Use Committee of the University of Pittsburgh (IACUC protocol number: 19126346). Two four-week-old female athymic nude (nu/nu) (NIH-Foxn1^rnu^) rats, purchased from Charles Rivers Laboratories, were maintained in a temperature-controlled animal facility at the UPMC Children’s Hospital of Pittsburgh with a 12-hour light/dark cycle. Animals were kept in the facility for at least two days prior to performing any procedures.

### Tumor grafts

Tumors were grafted as previously described in mice, but in four-week-old RNU homozygous rats (Charles Rivers Laboratories).(11) Under isoflurane anesthesia, a transverse left flank incision was made; and the left kidney and adrenal gland were exteriorized, then maneuvered using a cotton tip applicator. Five million SK-N-BE (2) neuroblastoma cells were suspended in 30 µL of Matrigel basement membrane matrix (Corning™ Cat#354234) and implanted into the left adrenal gland and surrounding fat pad with a 23-gauge needle on a 1cc syringe and the needle was slowly removed. The kidney and adrenal gland were returned to the abdominal cavity, the body wall was closed with 4-0 Polysorb suture, and the skin was closed with surgical wound clips. The animals recovered in a clean cage on a heating pad and were monitored until sternally recumbent. Wound clips were removed 10-14 days later.

### Surgery

Tumor resections were performed at 48 (rat 1) and 49 days (rat 2) after SK-N-BE (2) injections. Under isoflurane anesthesia, the rat was laid supine, and a lateral transverse abdominal incision was made (Fig 3A). Sharp dissection combined with a fine-tip cautery tool (BVI Accu Temp) was used to open the abdominal wall and meticulously separate the tumor from the surrounding healthy tissue. The adrenal vein was ligated with GEM micro clips (GEM, Synovis) or suture ties then transected while smaller blood vessels were cauterized. The left kidney was removed with the specimen if it could not be safely isolated. Following resection, tumor were sent for pathologic analysis.

### Pathology

Tumor samples were fixed in 10% neutral buffered formalin and processed and stained with hematoxylin and eosin according to standard protocols. (12) Formalin-fixed, paraffin-embedded tissue sections were stained on the Ventana BenchMark Ultra automated staining platform. Slides were pretreated with ultraCC1 (proprietary, Roche Tissue Diagnostics, Indianapolis, IN) and stained using a recombinant antibody to PHOX2B (EPR14423) (Abcam, Cambridge, MA, ab183741). OptiView DAB detection kit (proprietary, indirect, biotin free system, Roche Tissue Diagnostics, Indianapolis, IN) was used for primary antibody detection. All slides were counterstained with hematoxylin and routinely dehydrated, cleared, and cover-slipped in resinous mounting media. Four slides per rat were assessed by both H&E staining and PHOX2B staining.

## Results

Here, we demonstrate a novel SK-N-BE (2) human neuroblastoma cell xenograft in the adrenal gland of RNU homozygous rats. Cells suspended in Matrigel were injected directly into the adrenal glands and surrounding fat pads of 4-week-old nude rats (Fig 1). The rats continued to grow, from 100 g initially, to 173 g and 168 g respectively 4 weeks after injection and peaked at 187 g and 193 g at 6 weeks after injection. At 4 weeks, the tumors were visible and palpable in the left flank of the rats (Fig 2 A i-iv). MRI scans at 5 weeks post injections revealed tumors of over 2 cm in both rats (1.74 × 3.11 cm and 1.70 × 2.15 cm, respectively), covering the ipsilateral kidney (Fig 2 B, i-iii).

**Figure 1.**
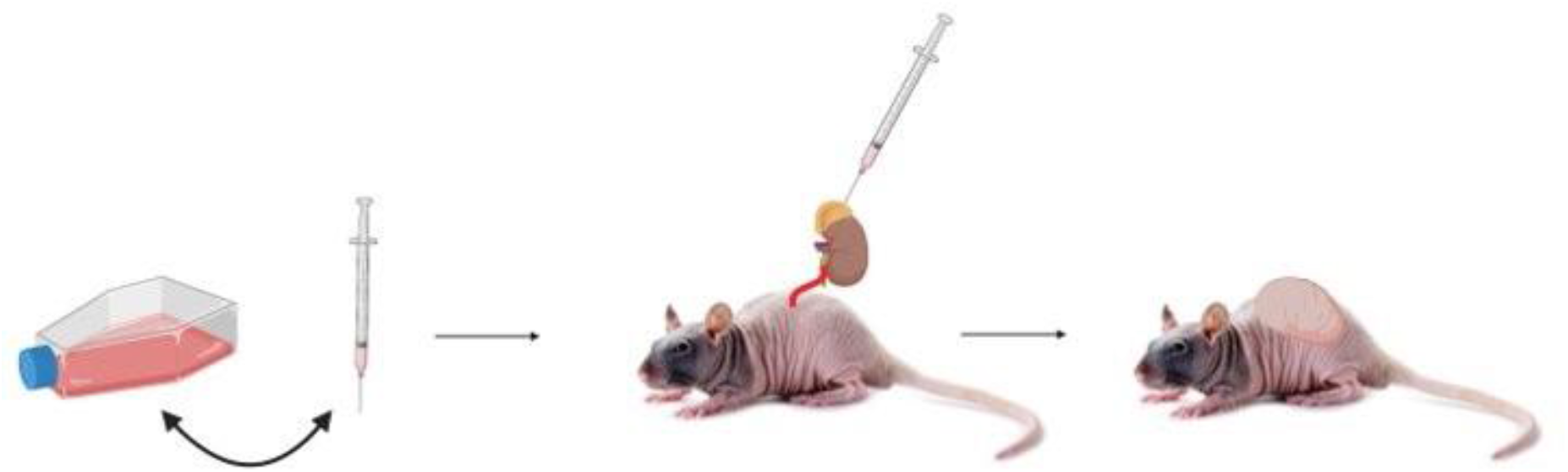
Schematic of orthotopic injection of SK-N-BE (2) cells into the left adrenal gland of 4-week old RNU rats (Created with BioRender.com).

**Figure 2.**
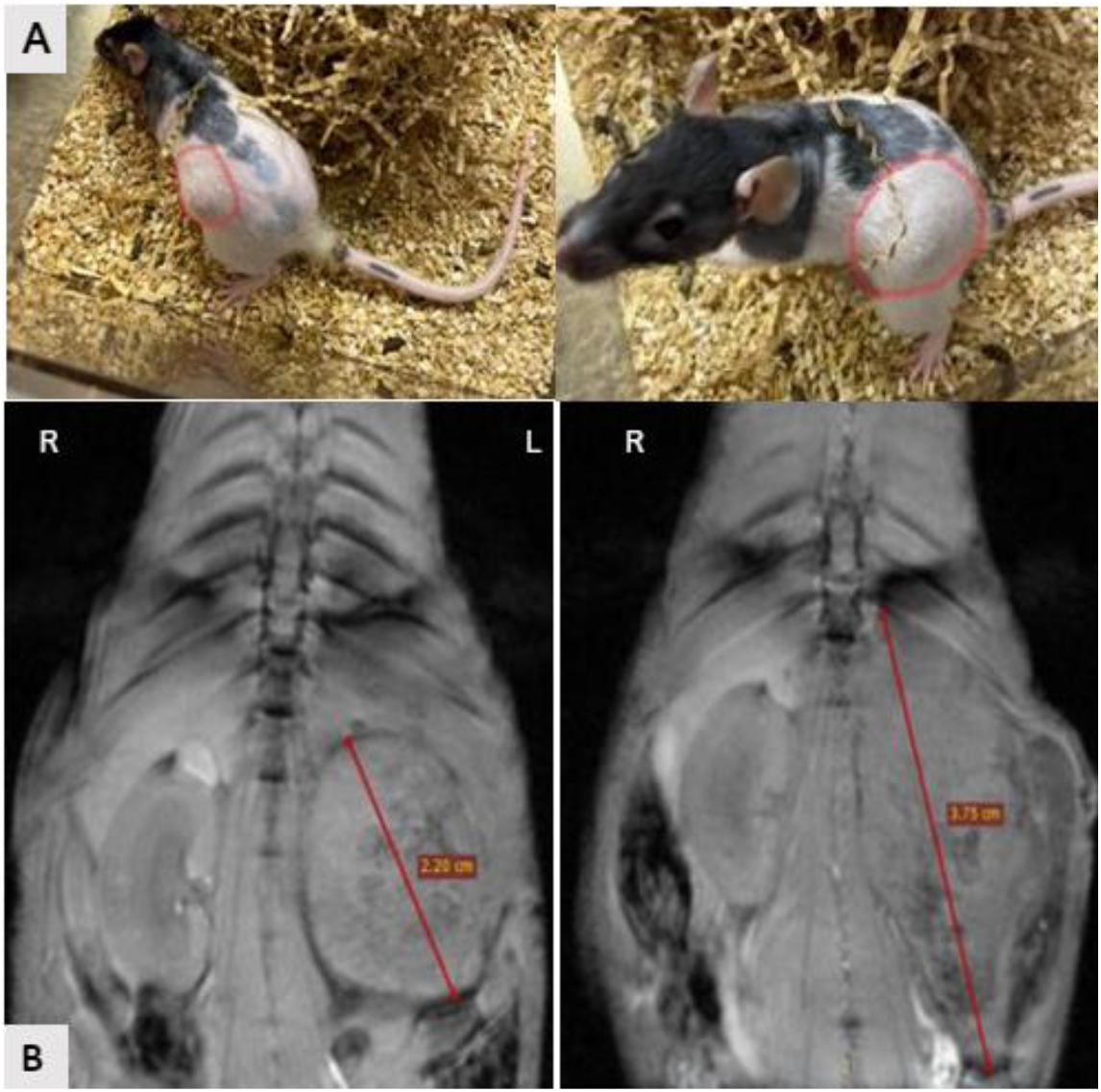
(A) White light images (rat 1 i, iii- rat 2 ii, iv) and (B) MRI images of neuroblastoma in the left adrenal gland of RNU rats 5 weeks post injection (i-coronal, ii- sagittal, iii-axial).

The rats began to lose weight at 7.5 weeks. They both exhibited hematuria which resolved. Rat 1 (Figure 3 A and B) continued to lose weight, up to 15% of its peak weight (187 g to 158.1 g), and surgery was attempted. The tumor had encased the left kidney and had adhesions to the stomach, spleen, and ureter. While attempting to separate the spleen from the tumor, the rat passed away. There had been no significant operative blood loss, so the death was most likely due to a combination of the stress of the large tumor as well as the stress of anesthesia and surgery. The tumor, which measured 4.1 × 4.5 × 1.5 cm was removed post-mortem and sent for pathology.

**Figure 3.**
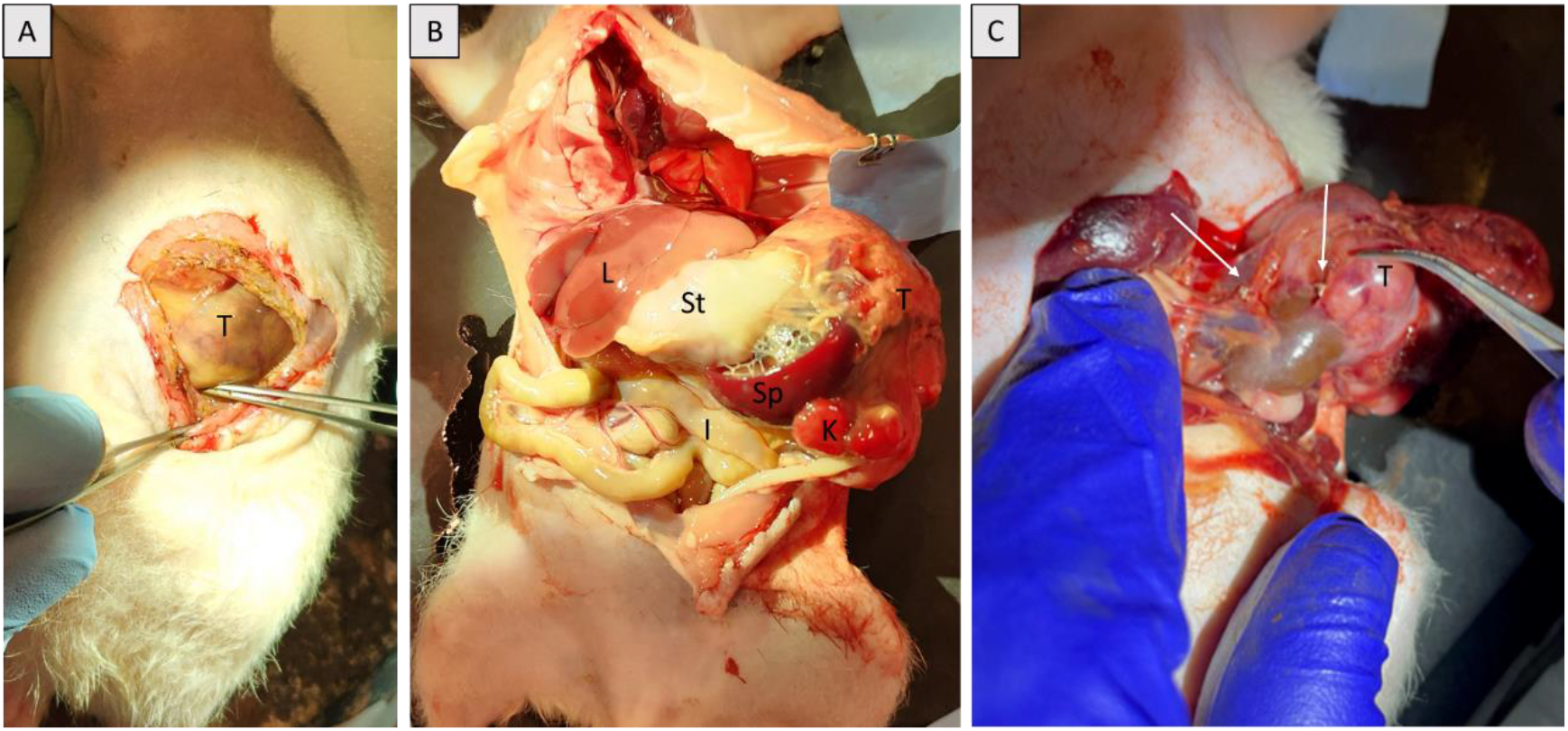
**(A)** Surgical incision for left adrenalectomy. **(B)** Exposure of tumor (T) in relation to abdominal organs including the liver (L), stomach (St), spleen (Sp), left kidney (K), and intestines (I). **(C)** Ligation of the adrenal vein using micro clips (arrows) (GEM, Synovis).

**Figure 4.**
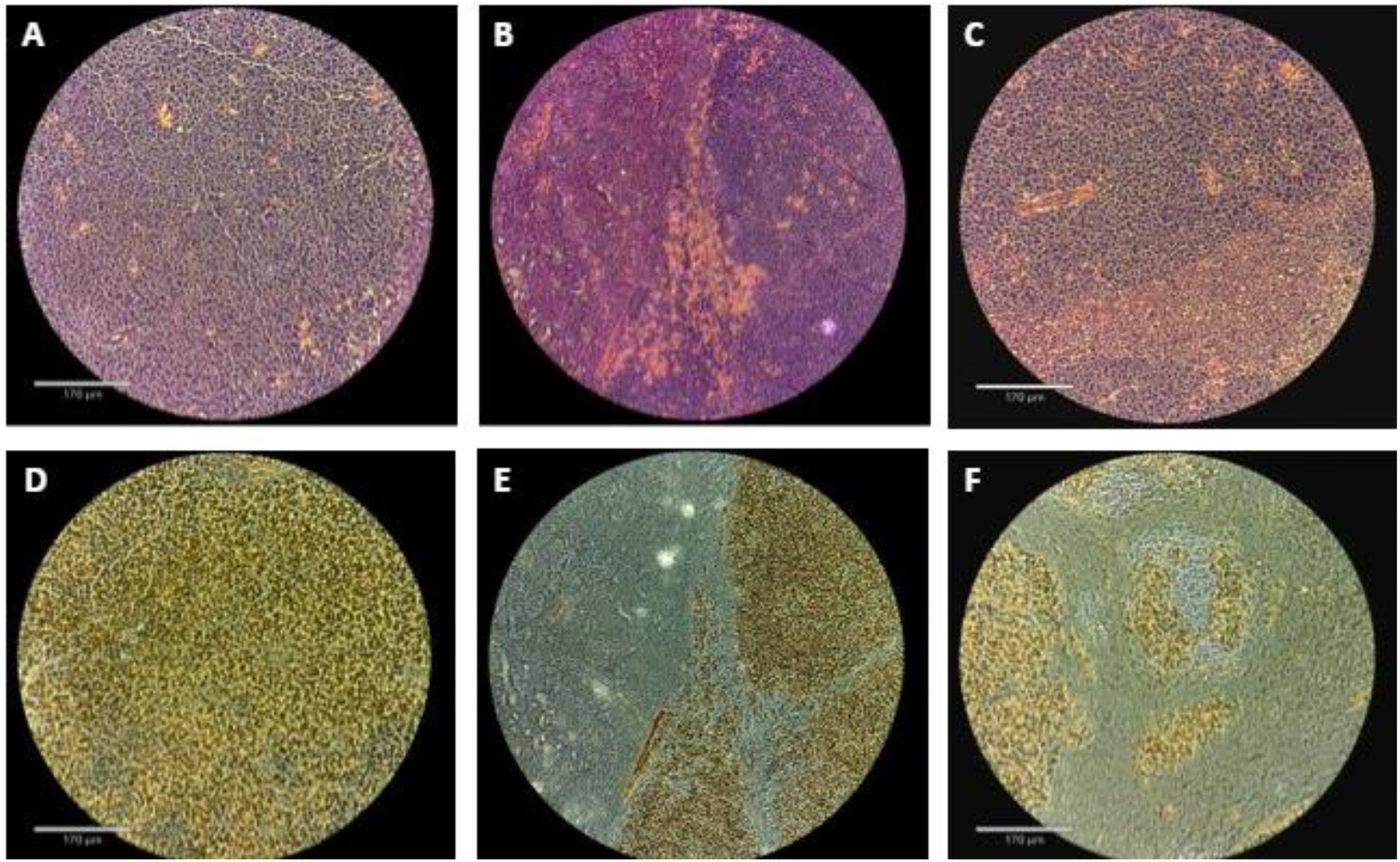
(Top) H&E staining demonstrating (A) small round blue cell tumor with Homer Wright pseudorosettes (rat 1), (B) same with invasion of renal parenchyma seen on the left (rat 2), and (C) with an area of hemorrhage. (Bottom) PHOX2B staining demonstrating (D) strong staining (rat 1), same with kidney invasion (E) (rat 2), and (F) an area of hemorrhage and necrosis.

Rat 2 lost about 5% of its total weight (193 g to 183.6 g) on the day of surgery (day 49). The tumor was smaller than that in rat 1 but also encased the left kidney and surrounding renal vessels. The adrenal vein ligation, a critical portion of surgery in humans, was successful using GEM micro clips (GEM, Synovis) (Figure 3 C). Unfortunately, this rat also passed away shortly after the adrenal vein ligation, again after minimal blood loss. The tumor measured 4.2 × 3.3 × 2.2 cm.

Pathology of both rats demonstrated poorly differentiated neuroblastoma with unfavorable histology. Small round blue cell tumors with Homer-Wright pseudorosettes and strong intranuclear PHOX2B staining was seen. There were large areas of hemorrhage and necrosis as well as focal invasion of adipose tissue and through the renal capsule in both tumors. Furthermore, both had a high mitosis karyorrhexis index, indicating unfavorable histology (Figure 5).

## Discussion

Despite multimodal therapy that includes surgical resection, high risk neuroblastoma in children remains fatal in 40-50% of patients within 5 years. (1,13) Safe and complete resection remains a challenge, largely due to encased neurovascular structures, difficult to detect tumor margins, and remote nodal metastases. (4) Unfortunately, completeness of resection likely impacts both patient outcomes and response to consolidation therapy with Dinutuximab. (14–16) Innovations in surgery are challenging to study with current rodent models, which are predominantly too small to replicate human disease (in mice) or are anatomically distinct from typical human disease (subcutaneous or tail vein injection-based models). The only described rat adrenal xenograft model was last published over 20 years ago. (8)

Here, we have developed a novel rat model of adrenal neuroblastoma using orthotopic injection of a human neuroblastoma cell line, SK-N-BE(2), which is well-established in its ability to form orthotopic tumors and widely utilized in preclinical studies. (refs) Furthermore, derivatives of SK-N-BE(2) cells have been shown to have amplified MYC-N, which is a feature of high-risk disease in humans. (3,17) Cells suspended in Matrigel were successfully injected directly into the adrenal glands and surrounding fat pads of 4-week-old nude rats (Fig 1). Using immunodeficient rodents is common in xenograft models to enable the growth of human tumor cell lines. The method of xenograft implantation is well described in mice and here has been translated to rats.

Tumors grew quickly and pathology was consistent with a poorly differentiated neuroblastoma with unfavorable histology, demonstrating small round blue cell tumors with Homer-Wright pseudorosettes. Both tumors also highly expressed PHOX2B, which is a sensitive and specific (versus other pediatric small round blue cell tumors) marker of neuroblastic tumors. (18) There were also large areas of hemorrhage and necrosis with focal invasion of adipose tissue and kidneys. Both also had a high mitosis karyorrhexis index, indicating unfavorable histology. While risk stratification in humans incorporates clinical features such as age, which cannot be truly replicated in models, poorly differentiated cancer with unfavorable histology likely recapitulates aggressive disease, which would benefit most from improved therapy. Cumulatively, the radiographic and histopathologic features are similar to those seen in humans.

By 4 weeks after xenograft implantation (8 weeks of age), tumors were of sufficient size for surgical resection. The rats began to lose weight at 6 weeks, however, suggesting excessive disease burden. Hematuria occurred, likely due to renal invasion. Surgery was attempted at 7 weeks, by which time the tumors had visually begun invading adjacent organs and the rats were unable to survive surgical intervention.

We intend to attempt surgical resection of these adrenal xenografts in the near future. We will perform surgery at an earlier timepoint, such as week 4, to allow the animals sufficient reserve to survive the stress of anesthesia and surgery. We were able to perform resection with minimal blood loss or trauma to adjacent organs, and believe surgery in a healthier animal can be performed with animal survival, as has been successfully achieved in mice (refs and unpublished data). We plan to assess recurrence of tumor and lifespan of the rats after surgical resection with or without the use of adjuncts, such as the use of intraoperative contrast agents. The increased size of the abdominal organs and tumor, as well as the adrenal localization of the xenograft in this novel rat model is critical for enabling these future studies.

## Conclusions (optional)

## Notes

### Competing Interest Statement

The authors have declared no competing interest.

